# Systematic characterization of chromodomain proteins reveals an H3K9me1/2 reader regulating aging in *C. elegans*

**DOI:** 10.1101/2022.08.10.503448

**Authors:** Xinhao Hou, Mingjing Xu, Chengming Zhu, Jianing Gao, Meili Li, Xiangyang Chen, Cheng Sun, Björn Nashan, Jianye Zang, Shouhong Guang, Xuezhu Feng

**Affiliations:** The USTC RNA Institute, Ministry of Education Key Laboratory for Membraneless Organelles & Cellular Dynamics, Department of Obstetrics and Gynecology, The First Affiliated Hospital of USTC, School of Life Sciences, Division of Life Sciences and Medicine, Biomedical Sciences and Health Laboratory of Anhui Province, University of Science and Technology of China, Hefei, Anhui 230027, China; CAS Center for Excellence in Molecular Cell Science, Chinese Academy of Sciences, Hefei, Anhui 230027, P.R. China

## Abstract

The chromatin organization modifier domain (chromodomain) is an evolutionally conserved motif across eukaryotic species. The chromodomain mainly functions as a histone methyl-lysine reader to modulate gene expression, chromatin spatial conformation and genome stability. Mutations or aberrant expression of chromodomain proteins can result in cancer and other human diseases. Here, we systematically tagged chromodomain proteins with green fluorescent protein (GFP) using CRISPR/Cas9 technology in *C. elegans*. By combining ChIP-seq analysis and imaging, we delineated a comprehensive expression and functional map of chromodomain proteins. We then conducted a candidate-based RNAi screening and identified factors that regulate the expression and subcellular localization of the chromodomain proteins. Specifically, we revealed a new H3K9me1/2 reader, CEC-5, both by in vitro biochemistry and in vivo ChIP assays. MET-2, an H3K9me1/2 writer, is required for CEC-5 association with heterochromatin. Both MET-2 and CEC-5 are required for the normal lifespan of *C. elegans*. Furthermore, a forward genetic screening identified a conserved Arginine124 of CEC-5’s chromodomain, which was essential for CEC-5’s association with chromatin and life span regulation. Thus, our work will serve as a reference to explore chromodomain functions and regulation in *C. elegans* and allow potential applications in aging-related human diseases.

## Introduction

The eukaryotic genome is packaged with histones and other proteins to form chromatin. Histones are subject to many types of posttranslational modifications (PTMs), especially on their flexible tails. These modifications include acetylation and methylation of lysine (K) and arginine (R) and phosphorylation of serine (S) residues, and play fundamental roles in most biological processes that are involved in chromatin dynamics, gene expression regulation and other DNA processes, such as repair, replication and recombination. To interpret PTMs of histones, effectors/readers are recruited to provide a link between the chromatin landscape and functional outcomes (Cui and Han 2007; Taverna *et al*. 2007; Bannister and Kouzarides 2011; Yun *et al*. 2011; Lawrence *et al*. 2016; DasGupta *et al*. 2020).

Over decades, multiple families of conserved domains that recognize modified histones have been discovered. These domains include members of the structurally related “Royal family,” such as chromo domains, tudor, PWWP and MBT (malignant brain tumor) repeat domains (Maurer-Stroh *et al*. 2003), which recognize mono-, di- or trimethylated lysine residues. The chromodomain is an evolutionally conserved region of approximately 30-60 amino acids (Lomberk *et al*. 2006). It was first identified in polycomb (Pc) proteins and heterochromatin protein 1 (HP1) (DasGupta *et al*. 2020). Despite the nature of recognition of methyl-lysine of histones (Taverna *et al*. 2007; Hyun *et al*. 2017), the biological functions of chromodomain proteins are highly diverse (Taverna *et al*. 2007; Hyun *et al*. 2017; DasGupta *et al*. 2020). For example, the (Pc) proteins and HP1 play important roles in maintaining facultative and constitutive repressive heterochromatin, respectively, through their recognition of methyl-lysine residues on histone H3 (H3K27me and H3K9me) (Taverna *et al*. 2007; Eissenberg 2012; DasGupta *et al*. 2020). In contrast, CHD1 (chromo-ATPase/helicase DNA-binding protein 1) was implicated in transcriptionally active regions in chromosomes. Furthermore, the double chromodomains of CHD1 cooperate with each other to bind to methylated H3K4 (Flanagan *et al*. 2005; Taverna *et al*. 2007). The spectrum of different chromodomain proteins may display a layer of regulation on the chromatin landscape, genome organization and gene expression. Thus, systematically delineating the complicated functional network of chromodomain proteins may provide further understanding of how and why these chromatin regulators function.

*C. elegans* has a number of beneficial features, making it a particularly powerful system for advancing our knowledge of chromodomain proteins and their functions in genome biology. The genome of *C. elegans* is compact, consisting of 100 Mb of DNA containing approximately 20,000 protein-coding genes. Importantly, due to the holocentric nature of *C. elegans* chromosomes, markers of heterochromatin, such as H3K9 methylation, are not concentrated at a single region on each chromosome. Instead, H3K9 methylation is enriched on chromosome arms in dispersed small domains (Liu *et al*. 2011). This enables easy sequencing of repetitive sequences. Moreover, endogenous fluorescent tagging coupled with microscopy, functional omics and rapid genetic screening makes it possible to efficiently assess the biological function of chromodomain proteins. Furthermore, the well-studied development, aging, stress response and short life cycle of *C. elegans* enable us to investigate the diverse roles of chromodomain proteins in physiological processes.

The *C. elegans* genome encodes 21 proteins that contain chromodomains, 9 of which have identified homologs in humans, yet most of the chromodomain proteins remain poorly characterized. The two *C. elegans* HP1 homologs (HPL-1 and HPL-2) physically associate with transcriptional repressive heterochromatin. HPL-1 has been found in an LSD-1/CoRESTlike complex (lysine-specific demethylase-1, corepressor for REST) (Vandamme *et al*. 2015; DasGupta *et al*. 2020). HPL-2 interacts with the zinc-finger protein LIN-13 and the H3K9me-binding MBT domain protein LIN-61, forming a complex that is part of the synthetic multivulva (synMuv) B group (Coustham *et al*. 2006; Harrison *et al*. 2007; Koester-Eiserfunke and Fischle 2011; Wu *et al*. 2012). CEC-1 and CEC-6 are two readers of H3K27me. CEC-1, together with the H3K9me reader CEC-3, contributes to the robust development, normal lifespan and fitness of animals. CEC-3 and CEC-6 are required for germline immortality maintenance (Saltzman *et al*. 2018). HERI-1 (also termed CEC-9) has been reported to antagonize nuclear RNAi by limiting H3K9me3 at siRNA-targeted genomic loci (Perales *et al*. 2018). UAD-2 is a newly identified chromodomain protein that recognizes H3K27me3 and promotes piRNA focus formation and transcription (Huang *et al*. 2021). It colocalizes with the upstream sequence transcription complex (USTC) at inner nuclear membrane (INM) foci (Weng *et al*. 2019) (Huang *et al*. 2021).

Susan Gasser and colleagues introduced a GFP reporter system using lacI/lacO repetitive arrays bearing the heterochromatic histone modifications H3K9me3 and H3K27me3 into *C. elegans* (Bessler *et al*. 2010; Meister *et al*. 2010). Genetic screens based on the subcellular localization of the GFP arrays identified regulators of heterochromatic formation and chromatin positioning at the nuclear periphery, including histone methyl transferases (HMTs) MET-2 and SET-25 and the chromodomain proteins CEC-4 and MRG-1 (Towbin *et al*. 2012; Gonzalez-Sandoval *et al*. 2015; Cabianca *et al*. 2019). In embryos, CEC-4 bound to the inner nuclear membrane and tethered heterochromatin through H3K9me to the nuclear periphery (Gonzalez-Sandoval *et al*. 2015; Harr *et al*. 2016). In intestinal cells, MRG-1 binds to euchromatin (H3K36me) and acts indirectly to anchor heterochromatin to the inner nuclear membrane (Cabianca *et al*. 2019).

Here, by systematically fluorescence tagging chromodomain proteins followed by ChIP-seq, genetic screening, and in vitro biochemical assays, we generated a resource of chromodomain proteins in *C. elegans*. The resource provides a comprehensive expression, regulation and functional map of the proteins in a eukaryotic system, which will not only serve as a reference to explore chromodomain proteins in *C. elegans* but also allow potential application in aging-related human diseases.

## Results

### A resource of fluorescence-tagged chromodomain proteins in *C. elegans*

The chromodomain is highly conserved in a wide range of organisms from *Schizosaccharomyces pombe, Drosophila melanogaster, Arabidopsis thaliana*, and *Caenorhabditis elegans* to *Mus musculus* and *Homo. sapiens*. According to the presence of other types of domains, chromodomain proteins can be classified into 13 families (Tajul-Arifin *et al*. 2003; DasGupta *et al*. 2020). These families include the chromodomain-helicase DNA-binding (CHD) family, the histone methyl transferase family, the HP1 family, the Polycomb family, the Msl-3 homolog family, the histone acetyltransferase (HAT) family, the retinoblastoma-binding protein 1 (RBBP1) family, the enoylCoA hydratase family, the SWI3 family, and the plant-specific chromomethylase family (Tajul-Arifin *et al*. 2003).

The *C. elegans* genome encodes 21 proteins that contain chromodomains, 9 of which have been reported to have homologs in humans (Table S1). The homologs are divided into 4 conserved families, including the CHD family, HP1 family, histone acetyltransferase family, and Msl-3 homolog family. CHD family members contain paired tandem chromodomains and helicase domains. Among them, CHD-1 and CHD-7 are homologs of human CHD1 and CHD5, respectively. CHD-3 and LET-418 are two Mi-2 homologs. The HP1 family members HPL-1 and HPL-2 are also conserved. Mortality factor-related gene (MRG-1) is the ortholog of mammalian MRG15. The two histone acetyltransferases MYS-1 and MYS-2 are conserved in humans as well (Tajul-Arifin *et al*. 2003). We performed sequence alignment and phylogenetic analyses of chromodomain proteins in *C. elegans* and *H. sapiens*. This result was largely consistent with previous work (Fig. 1A). Notably, we identified CEC-7 as a homolog of the Msl-3 family proteins of humans (Fig. 1A). Although most *C. elegans* chromodomain proteins (*cec* genes) have no clear homologs in humans, the chromodomains of these genes share high identity with the chromodomain of human HP1α (Table S2).

**Figure 1.**
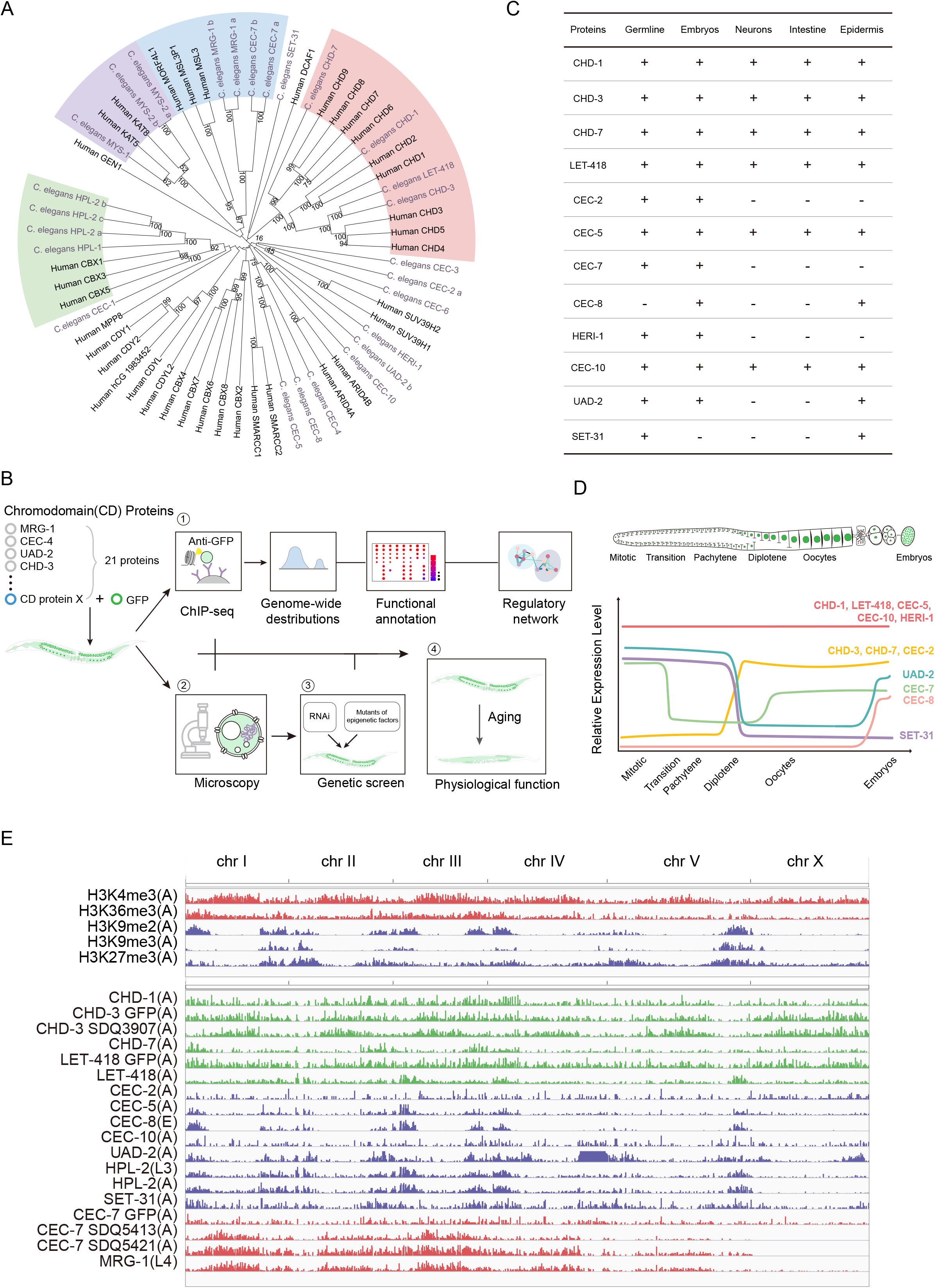
Systematic analysis of chromodomain proteins in *C. elegans*. (A) Phylogenetic tree of chromodomain proteins in *C. elegans* and *H. sapiens*. (B) Schematic diagram for systematic analysis of chromodomain proteins. See text for details. (C) Summary of the expression patterns of chromodomain proteins in the indicated cells. (D) Curve graph showing the expression profiles of chromodomain proteins in the germline. (E) Distribution of ChIP-seq peaks of active, repressive chromatin marks (upper panel), and chromodomain proteins (lower panel) across the genome. The development stage of animals for each sample is noted as E (embryos), L (larval), and A (adult).

To investigate the expression and function of these chromodomain proteins, we used CRISPR/Cas9 technology and constructed a number of fluorescently tagged transgenic strains by knocking in a *gfp-3xflag* tag in situ to the chromodomain genes. These strains were subjected to assays, including ChIP-seq, fluorescence microscopy, and physiological function annotation. By systematically analyzing the ChIP-seq datasets, we delineated the genome-wide distribution, functional pathways, and epigenetic regulatory network in which these chromodomain proteins participate. In addition, we used the subcellular localization of each chromodomain protein as a reporter and screened for genetic factors that regulate the expression and localization of chromodomain proteins. The experimental pipeline is shown in Fig. 1B.

Of the 21 chromodomain genes in *C. elegans*, we successfully targeted 11 of them with a GFP-3xFLAG fluorescent tag. Nearly all of the chromodomain proteins were broadly expressed in nuclei throughout germline, somatic cells and embryos (Figs. 1C and S1A-B). Interestingly, the expression of many chromodomain proteins was dynamically altered in the germline (Figs. 1D and S1A). For example, UAD-2 and SET-31 were expressed in the mitotic and meiotic regions but not in oocytes (Huang *et al*. 2021). In contrast, CHD-3, CHD-7, and CEC-2 were not expressed in the mitotic and early meiotic regions but began to be expressed in diplotene cells and oocytes. Notably, CEC-7 exhibited a dramatic reduction in expression at the transition zone but was re-expressed in oocytes (Figs. 1D, S1A). CEC-8 was not expressed in the germline. Knockout of these chromodomain proteins slightly reduced the brood size of animals (Fig. S1C). Together, these data implied an orchestrated regulation of chromodomain proteins during germ cell maturation.

To identify genome binding signatures of the chromodomain proteins, we used GFP antibody (#ab290) and conducted ChIP-seq experiments in adults or late embryos from the fluorescently tagged strains (Fig. 1E and Table S3). In addition, reported ChIP-seq datasets of several chromodomain proteins were downloaded from the NCBI GEO database (Table S3). *C. elegans* chromosomes are organized into broad domains that differentiate the center of the chromosomes from the arms: active chromatin marks such as H3K4me3 and H3K36me3 have a similar distribution from centers to arms, while repressive histone marks, especially H3K9me1/2/3, are enriched at the distal chromosome arms (Liu *et al*. 2011). We mapped the binding locations of chromodomain proteins and compared the patterns to each other and to those of H3K9me2, H3K9me3, H3K27me3, H3K4me3, and H3K36me3 marks. Remarkably, approximately 44% of gene sequences are localized on chromosome arms. CHD-7 (58% on arms), CEC-5 (66% on arms), CEC-8 (97% on arms), CEC-10 (59% on arms), UAD-2 (77% on arms), HPL-2 (71% on arms in young adult; 68% on arms in L3), and SET-31 (62% on arms) were preferentially enriched at chromosome arms of autosomes. CHD-3 (35% on arms), CHD-1 (45% on arms), LET-418 (43% on arms in this study), CEC-2 (36% on arms), MRG-1 (32% on arms), and CEC-7 (46% on arms) were uniformly distributed from chromosome arms to centers (Figs. 1E, S2A). The diverse genome distribution patterns implied distinct functions of chromodomain proteins in chromatin regulation.

### Association maps between chromodomain proteins and histone modifications on chromosome arms and centers

Chromodomain proteins typically bind specific histone modifications (Taverna *et al*. 2007; DasGupta *et al*. 2020). For example, CEC-4 recognizes H3K9me1/2/3, whereas MRG-1 may associate with H3K36me2/3 marks (Cabianca *et al*. 2019). To determine the histone modifications bound by each chromodomain protein in vivo, we analyzed the correlation of chromodomain protein distribution signatures with H3K9me2/3, H3K27me3, H3K4me3, H3K36me3 and H3K79me3 marks on chromosome arms and centers.

Chromodomain proteins displayed diverse propensities for histone modifications on chromosome arms that were enriched for both heterochromatin and euchromatin (Figs. 2A-E, S2B-E). Strikingly, CEC-8, HPL-2, and CEC-5 were prominently enriched in H3K9me2 abundant regions (Figs. 2A and 2E). A small portion of CEC-5 targets were coated with H3K4me3. UAD-2, CHD-7 and SET-31 exhibited similar distribution patterns and preferentially correlated with H3K27me3 marks, yet weak signals of H3K9me2/3 were also identified on the target sites (Figs. 2B and 2E). In contrast, CHD-3 and CHD-1 were strongly correlated with H3K4me3 marks (Figs. 2C and 2E). MRG-1 and CEC-7 were mainly correlated with H3K36me3 and H3K79me3, and CEC-7 might also associate with H3K9me2 and H3K4me3 (Figs. 2D-E, S2E).

**Figure 2.**
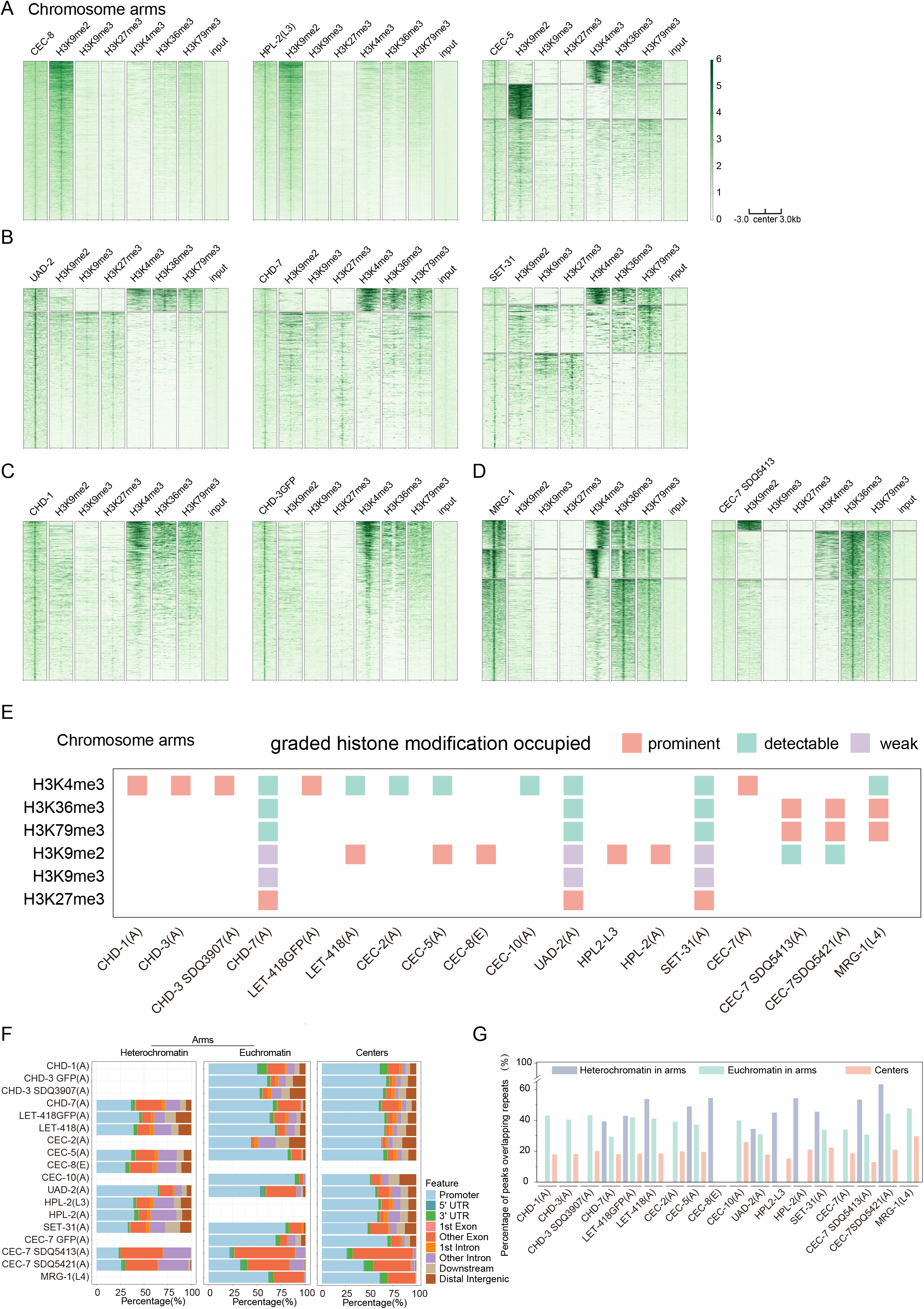
Distinct chromodomain protein-histone modification association maps on chromosome arms and centers. (A, B, C, D) Heatmap comparing heterochromatin-enriched histone modifications (H3K9me2, H3K9me3, H3K27me3) and euchromatin-enriched histone modifications (H3K4me3, H3K36me3, H3K79me3) with the indicated chromodomain proteins on chromosome arms. (E) A summary of the interaction of chromodomain proteins with histone modifications on chromosome arms. (F) Distribution of chromodomain protein ChIP-seq peaks on different genomic features (promoters, exons, introns, etc.) across distinct broad chromosome domains. (G) Bar plot showing the association of chromodomain proteins with repetitive elements across distinct broad chromosome domains on chromosomes.

In chromosome centers, which were enriched for active chromatin, most of the chromodomain protein targets, except MRG-1, were marked with H3K4me3 (Fig. S3A-3B). MRG-1 specifically correlated with H3K36me3 and H3K79me3 at the centers (Figs. S3A-3B). UAD-2, CHD-7 and SET-31 were correlated with H3K27me3, H3K36me3 and H3K79me3 (Figs. S3A-3B).

Most chromodomain proteins were associated with both heterochromatin and euchromatin (Fig. 2E) but exhibited different binding patterns in the two regions (Fig. 2F). In euchromatin, chromodomain proteins mainly bound to gene promoters, whereas in heterochromatin, chromodomain proteins targeted a large number of exons and introns (Fig. 2F). In addition, chromodomain proteins showed a greater propensity for repetitive sequences in heterochromatin regions (Fig. 2G).

Collectively, we revealed distinct chromodomain protein-histone modification association maps in chromosome arms and centers. Our data indicated the complicated function and mechanisms of the interaction of chromodomain proteins and histone modifications in vivo.

### Functional annotation of chromodomain proteins

To investigate the function of chromodomain proteins, we annotated the binding targets of each protein.

Most chromodomain proteins were enriched on protein coding genes (Fig. S4A). UAD-2 was highly enriched in piRNA genes, which was consistent with our previous work showing that UAD-2 mediates heterochromatin-directed piRNA expression (Fig. S4A) (Huang *et al*. 2021). Then, we performed pathway enrichment analysis of the gene targets and divided the pathways into “Development”, “Aging”, “Stress response”, “Cell cycle”, and “Biosynthesis & Metabolism” related terms (Fig. S4B). Most chromodomain proteins were enriched in a large number of biological pathways (Fig. S4B), suggesting general roles of the chromodomain proteins in these biological processes. UAD-2 was not enriched in any specific GO (Gene Ontology) terms, which is consistent with its preferential binding to piRNA genes other than protein coding genes (Fig. S4B) (Huang *et al*. 2021). CEC-8 and CEC-10 were also depleted from many specific GO (Gene Ontology) terms (Fig. S4B).

A portion of each chromodomain protein’s peaks overlapped with at least one repeat sequence (19.99-53.66%) (Fig. 2G). To test whether chromodomain proteins are biased toward specific repetitive sequence families, we classified 84,972 individual repetitive elements into 111 repeat families, which were further classified by sequence type (e.g., DNA transposon, retrotransposon, satellite, or unknown) (McMurchy *et al*. 2017). Among the 111 repeat families, forty-two were mostly DNA transposons, including the transposase and Helitron families, and were bound by at least one chromodomain protein (Fig. S5A).

### Epigenetic landscape of the chromodomain proteins

In eukaryotes, a plethora of histone modifications, writers, readers, erasers and remodelers cooperate to translate the chromatin epigenetic landscape into transcriptional activation or repression and genome stability modulation (Hyun *et al*. 2017).

To investigate the function of chromodomain proteins in the context of the epigenetic regulatory network, we downloaded previously published ChIP-seq datasets of a number of histone modification factors and epigenetic regulators. We first compared the correlation of the genome-wide distribution of these factors with that of the chromodomain proteins. We conducted hierarchical clustering of the genome-wide correlation coefficients and identified three major groups (Fig. 3A). The comparative analysis of all datasets was consistent with their putative functional relationships. Marks that are known to act in related pathways, such as transcriptional activation or repression, were highly correlated and clustered together. Group A contained repressive marks, including H3K9me3 and H3K27me3. Group B contained H3K9me2 and regulating factors such as MET-2, LIN-13, LIN-61 and HPL-2 (Zeller *et al*. 2016; McMurchy *et al*. 2017). LEM-2 (ceMAN1), a nuclear membrane protein associated with heterochromatin, was also included (Barkan *et al*. 2012). In addition, H3K9me1 in embryos, H3K36me1 in adults, and H3K27me1 in embryos were included in this group. Remarkably, most of the chromodomain proteins studied in this work (10 out of 13) were highly correlated with each other and clustered into Group B, suggesting similar functions of these chromodomain proteins on repressive chromatin. Group C was mainly composed of active histone marks and related proteins. Of these, H3K4me1/2/3 were clustered in the same subcluster, whereas H3K36me2/3 and H3K79me2/3 shared more similar profiles. The chromodomain proteins CHD-3 and LET-418 were clustered with H3K4me1/2/3. MRG-1 and CEC-7, two members of the Msl-3 family, were contained in the subgroup, including H3K36me2/3 and H3K79me2/3. The finding that MRG-1 was highly correlated with H3K36me2/3 was consistent with previous research showing that MRG-1 and its homologs in yeast and humans are deposited in euchromatic regions bearing H3K36me (Doyon *et al*. 2004; Cabianca *et al*. 2019).

**Figure 3.**
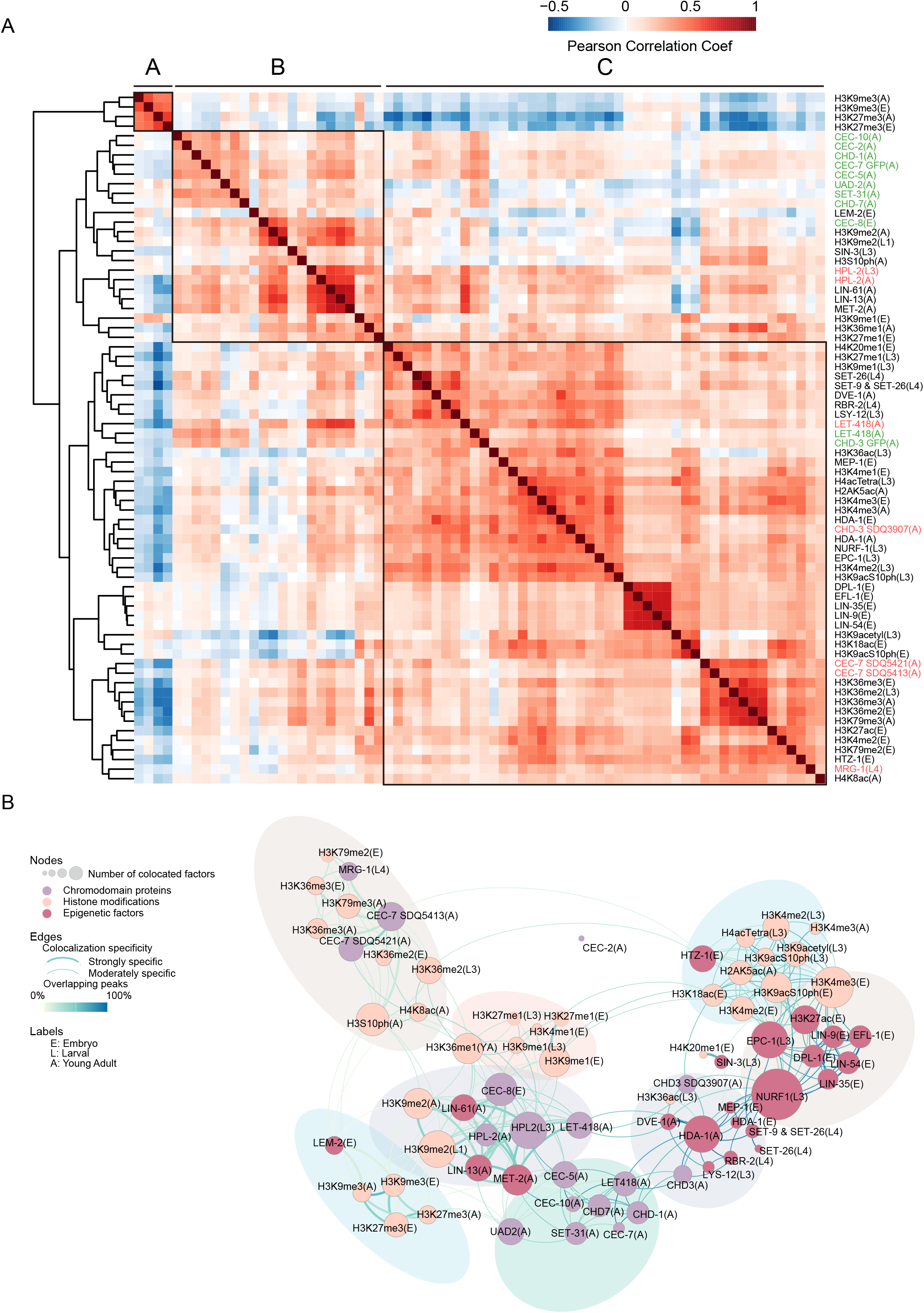
An epigenetic regulatory network of chromodomain proteins. (A) Hierarchical clustering of chromodomain proteins and epigenetic regulators based on the pairwise Pearson correlations of signals of ChIP-seq peaks (see Materials and methods for details). (Red) Positive correlations; (blue) negative correlations. The development stage of animals for each sample is noted as E (embryos), L (larval), and A (adult). (B) A colocalization network across the genome of chromodomain proteins and epigenetic factors. In the network, nodes indicate individual chromodomain proteins (purple), histone modifications (light pink) and other epigenetic regulators (dark pink); subnetworks identified are indicated by shadows in different colors; edge colors depict the percentages of overlap between the nodes and weight the colocalization specificity between two factors (see Materials and methods for details). The development stage of animals for each sample is noted as E (embryos), L (larval), and A (adult).

To further identify specific functional combinations or communities of the chromodomain proteins, histone modifications and their regulators, we generated a colocalization network based on the overlap of binding sites observed in the ChIP-seq datasets (Fig. 3B). We calculated the overlap significance between the binding sites of each pair of factors and the percentage of overlapping sites. We then identified strong and moderate colocalization significance of the factors (see Materials and Methods). Overall, we identified 8 communities in the network (Figs. 3B, S6A-C), which is similar to that of the hierarchical clustering approach. Most of the chromodomain proteins were highly interconnected with each other and exhibited a strong correlation with hallmarks of repressive chromatin, especially H3K9me2 and H3K27me3. CHD-3 and LET-418 displayed a high propensity for H3K4me1/2/3-related factors. MRG-1 and CEC-7 were present in the subnetwork containing H3K36me2/3 and H3K79me2/3. Detailed information on the epigenetic network is listed in Table S4.

Taken together, we delineated an epigenetic regulatory landscape involving chromodomain proteins by combining clustering and network analysis of a plethora of ChIP-seq datasets. Our analysis sheds light on the potential diverse roles and mechanisms of chromodomain proteins in epigenetic regulatory networks. The data will facilitate the elucidation of the biological roles and regulation of many uncharacterized chromodomain proteins.

### Distinct subcellular localization of the chromodomain proteins

We then profiled the subcellular localization of each chromodomain protein and assigned them to one or more of the 3 subcellular localization patterns: nucleoplasm, nucleolus, and nuclear punctate. Eight chromodomain proteins, CHD-1, CHD-3, CHD-7, LET-418, CEC-7, CEC-8, HERI-1 and CEC-10, were localized to the nucleoplasm (Fig. 4A). Interestingly, four heterochromatin-related proteins, CEC-5, CEC-8, SET-31, and UAD-2, accumulated in certain subnuclear compartments (Figs. 4B-C) (Huang *et al*. 2021). Our previous work showed that UAD-2, together with an upstream sequence transcription complex (USTC), colocalized with piRNA foci that were associated with piRNA clusters on chromosome IV (Weng *et al*. 2019; Huang *et al*. 2021).

**Figure 4.**
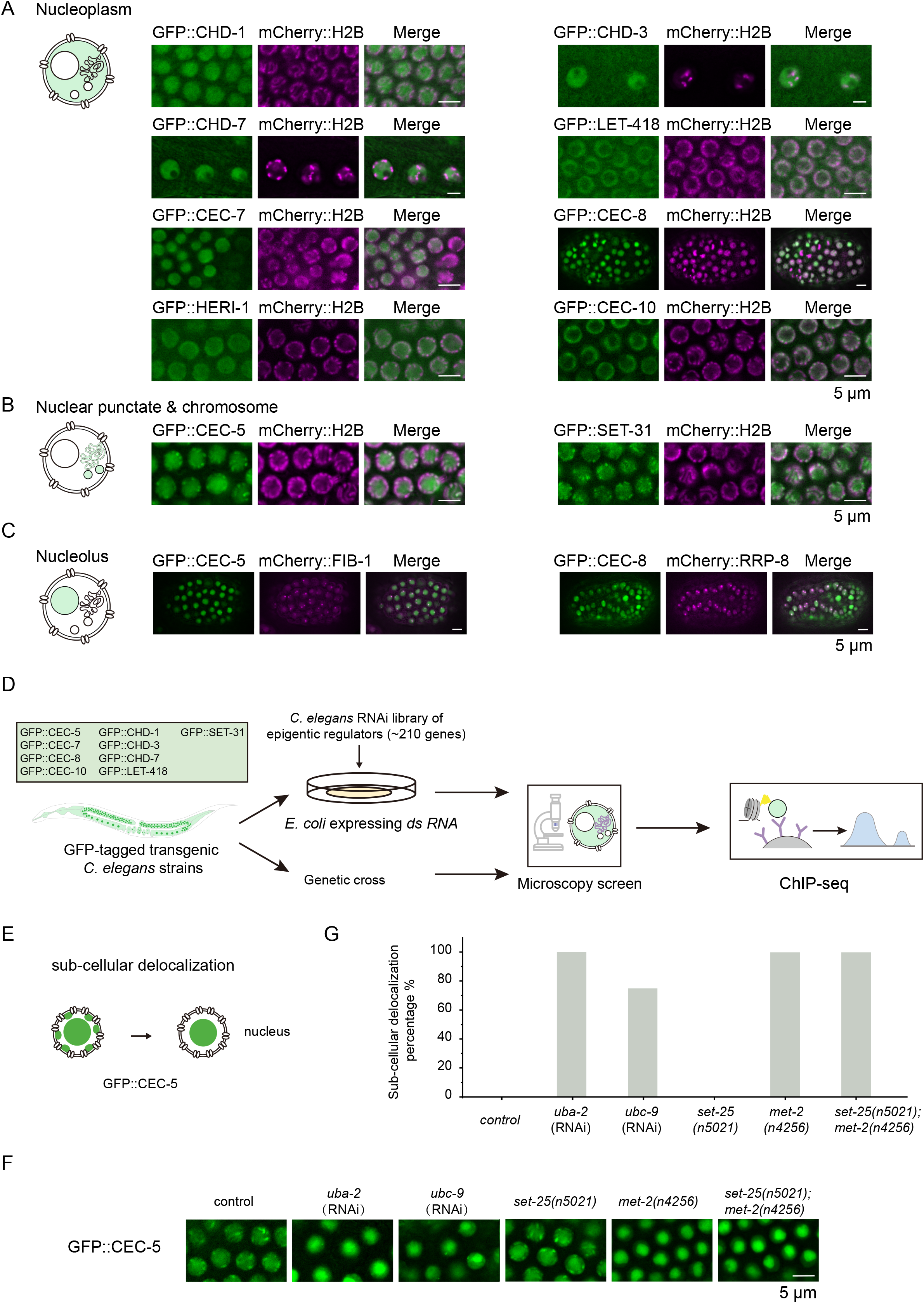
A candidate-based RNAi screening to search regulators of chromodomain proteins. (A, B, C) The 3 distinct subcellular localization patterns of chromodomain proteins; (A) Nucleoplasm in the germline; (B) Nuclear puncta and chromosomes in the germline; (C) Nucleolus in embryos. The localization of a representative image for each chromodomain protein is shown (scale bar, 5 μm). The chromosome is marked by mCherry::H2B, and the nucleolus is marked by mCherry::FIB-1 or mCherry::RRP-8. (D) Schematic procedure of the candidate-based RNAi screening. The subcellular localization of each chromodomain protein is used as a reporter. A library of 217 genes involved in chromatin modification and histone modification were selected to be knocked down or knocked out. (E) Schematic diagram of the screening for CEC-5 regulators. (F) Images of representative germline nuclei of the indicated adult animals. Knockdown of *uba-2* and *ubc-9* or mutation of *met-2* suppressed CEC-5 nuclear foci formation. (G) Quantification of the subcellular delocalization percentage in the indicated animals; N>50.

CEC-5 exhibits two localization patterns. CEC-5 was discontinuously distributed on chromosomes and was enriched in a number of nuclear puncta that were colocalized with mCherry::H2B in early embryos and germline (Figs. 4B, S7B). In addition, CEC-5 colocalized with the nucleolar marker mCherry::FIB-1 (Figs. 4C, S7C-D), which is consistent with the ChIP-seq data showing that CEC-5 predominantly occupied the 18S and 26S rDNA genes (Fig. S7E).

SET-31 was expressed in the germline and enriched on chromosomes. In mitotically proliferating cells, SET-31 colocalized with mCherry::H2B (Fig. S7B) and formed nuclear puncta in germ cells (Fig. 4B).

CEC-8 was also localized in the nucleolus. In a small proportion of embryo cells (∼15%), we observed that CEC-8 formed two distinct nuclear foci that colocalized with the nucleolus marker mCherry::RRP-8 (Figs. 4C, S7D). We did not detect significant binding of CEC-8 to either 18S or 26S rDNA genes in the ChIP-seq experiment (Fig. S7E).

The GFP-tagged transgenes could provide valuable reporter systems for investigating the function of chromodomain proteins, spatial distribution and condensation of heterochromatin and nucleolar regulation.

### A candidate-based RNAi screening to identify regulators of chromodomain proteins

Previously, we selected 239 genes involved in chromatin regulation and histone modification and designed a candidate-based RNAi screen to search for regulators of UAD-2 and piRNA transcription (Huang *et al*. 2021). We successfully identified a number of genes that are required for piRNA focus formation and piRNA transcription (Huang *et al*. 2021).

Here, to identify regulators of other chromodomain proteins, we used their subcellular localization as reporters to search for factors regulating the expression and localization of chromodomain proteins by fluorescence microscopy. Nine GFP-fused chromodomain proteins were investigated (Fig. 4D). We found that knocking down *uba-2 and ubc-9* and the mutation of *met-2* significantly inhibited the nuclear puncta formation of CEC-5 (Figs. 4E-G).

UBA-2 is an E1 protein, and UBC-9 is a single E2 SUMO-conjugating enzyme in *C. elegans* (Broday 2017). Modification by the small ubiquitin-related modifier (SUMO), known as SUMOylation, includes distinct enzymatic pathways that conjugate SUMO to target proteins. Recent studies have linked SUMOylation to several chromatin regulation processes (Cubenas-Potts and Matunis 2013). The observation that SUMOylation affects CEC-5 localization (Figs. 4F-G) suggested that SUMOylation may change the chromatin environment and modulate the function of CEC-5.

Strikingly, mutation of MET-2 disrupted the nuclear punctate formation of CEC-5 in the germline, whereas mutation of SET-25 did not affect CEC-5 localization (Figs. 4F-G). In *met-2* mutants, CEC-5 is significantly enriched in nucleoli. MET-2 is the homolog of mammalian SETDB1 that mediates mono- and dimethylation of H3K9. SET-25 deposits H3K9me3 marks in a MET-2-LIN-61- and NRDE-3-dependent manner (Padeken *et al*. 2021). These data suggested that CEC-5 may be regulated by H3K9me1/2.

To further assess the roles of MET-2 and H3K9me1/2 in the regulation of CEC-5, we performed ChIP-seq experiments of GFP::CEC-5, GFP::CEC-5;*met-2*, GFP::CEC-5;*set-25* and GFP::CEC-5;*met-2*;*set-25* in young adult animals. Notably, GFP::CEC-5 and GFP::CEC-5;*set-25* exhibited similar binding patterns; CEC-5 mainly bound chromosome arms and was associated with heterochromatin (Figs. 5A-C). However, in the *met-2* single mutant and *met-2*;*set-25* double mutant, CEC-5 significantly reduced its association with chromosome arms and heterochromatin, whereas the association of CEC-5 with chromosome centers and euchromatin was increased (Figs. 5A, 5D-E). We identified 134 coupregulated and 276 codownregulated CEC-5 targets that were enriched in both *met-2* single mutants and *met-2*;*set-25* double mutants (Figs. 5F, 5G, 5I). Interestingly, the coupregulated sites were uniformly distributed from chromosome centers to arms (Fig. 5H), whereas the codownregulated targets were mainly distributed on chromosome arms (Fig. 5J). Together, these results suggested that MET-2 and likely H3K9me1/2 were required for the proper association of CEC-5 with heterochromatin.

**Figure 5.**
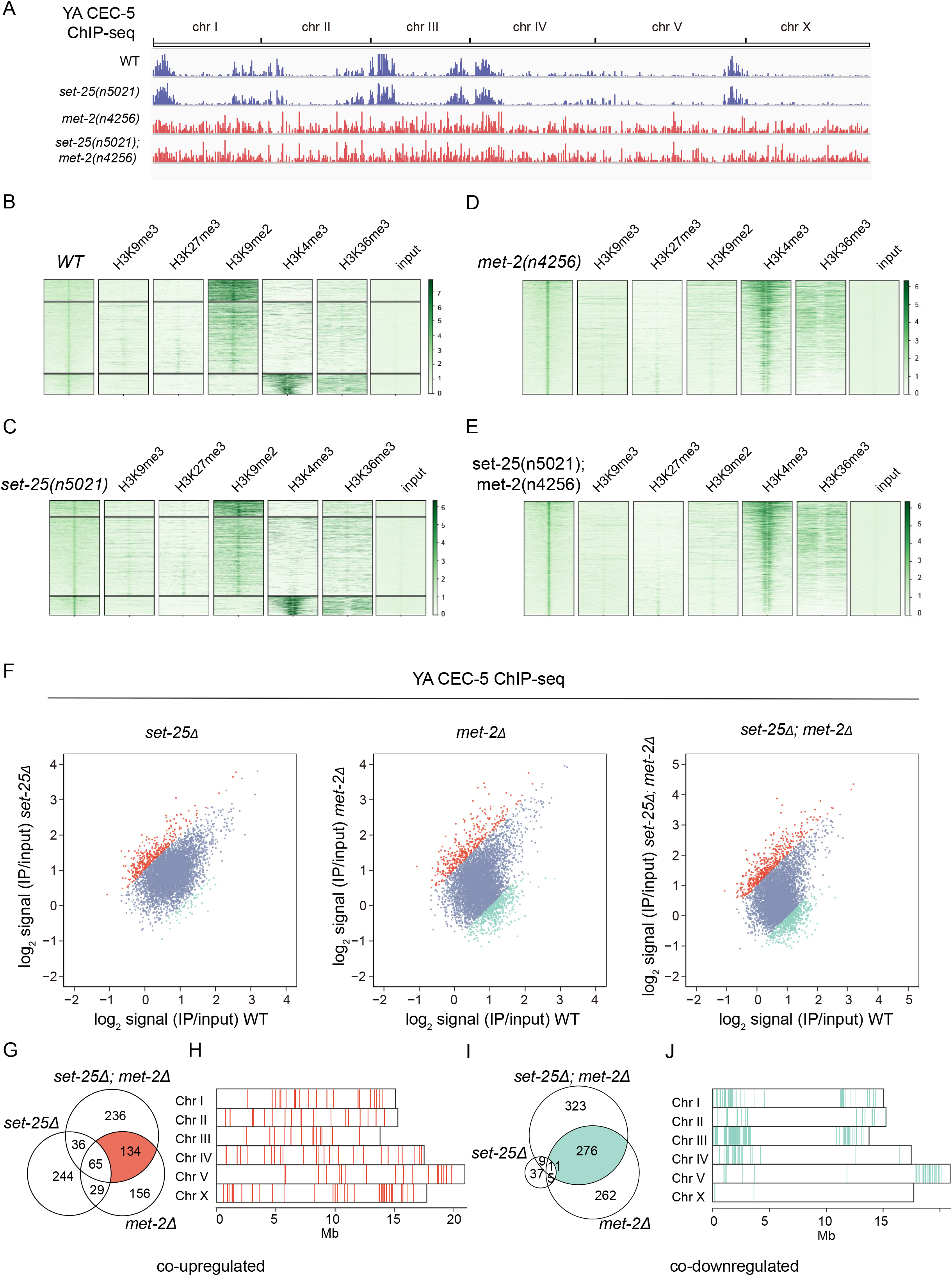
The association of CEC-5 with chromatin depends on H3K9me1/2. (A) ChIP-seq peak distribution of CEC-5 in the indicated adult animals. (B, C, D, E) Heatmap comparing heterochromatin histone modifications (H3K9me2, H3K9me3, H3K27me3) and euchromatin histone modifications (H3K4me3, H3K36me3, H3K79me3) with CEC-5 in the indicated animals. (F) Scatterplot comparing CEC-5 ChIP-seq signals in *set-25*Δ, *met-2*Δ, or *set-25*Δ; *met-2*Δ with wild-type animals. Differentially enriched peaks (>2-fold) for each genotype versus the wild type are highlighted in color (red, upregulated; blue, unchanged; green, downregulated). (G, H, I, J) Chromosome distribution of coupregulated peaks (G, H, red) and codownregulated peaks (I, J, green) in *met-2*Δ, and *set-25*Δ; *met-2*Δ compared to wild-type animals.

### The chromodomain of CEC-5 binds H3K9me0/1/2/3 in vitro

To assess whether CEC-5 directly binds to histones, we expressed and purified the GST-fused chromodomain-containing fragments of CEC-5 (CEC-5 CD, 51-172 aa). The fragments were incubated with biotin-labeled nonmethylated histone H3 peptides or methylated H3 peptides followed by precipitation with streptavidin agarose beads. The pelleted proteins were then resolved by SDS□PAGE and stained with Coomassie blue (Figs. 6A, 6B). As controls, GST and a GST-fused CEC-4 chromodomain-containing fragment (CEC-4 CD, 25-141 aa) were used (Gonzalez-Sandoval *et al*. 2015; Huang *et al*. 2021). The CEC-4 chromodomain has been shown to bind H3K9me1/2/3 in vitro (Gonzalez-Sandoval *et al*. 2015; Huang *et al*. 2021). Consistent with previous results, CEC-4 bound to H3K9me1/2/3 peptides (Figs. 6C, 6D) but not H3K9me0, H3K27me0, or H3K27me3 peptides (Figs. 6E, 6F). CEC-5 did not bind either H3K27me0 or H3K27me3 peptide (Figs. 6E, 6F). Surprisingly, CEC-5 bound both unmethylated (me0) and methylated (me1/2/3) H3K9 peptides, although CEC5 CD showed weaker binding affinity to unmethylated (me0) H3K9 peptide (Figs. 6C, 6D). Since the binding of CEC-5 to chromosomes depends on H3K9me1/2 but not H3K9me3 in vivo, we speculated that other amino acid sequences of CEC-5 may modulate the binding specificity of the CEC-5 chromodomain. Alternatively, certain cofactors of CEC-5 may help to determine the binding specificity of CEC-5 in vivo.

**Figure 6.**
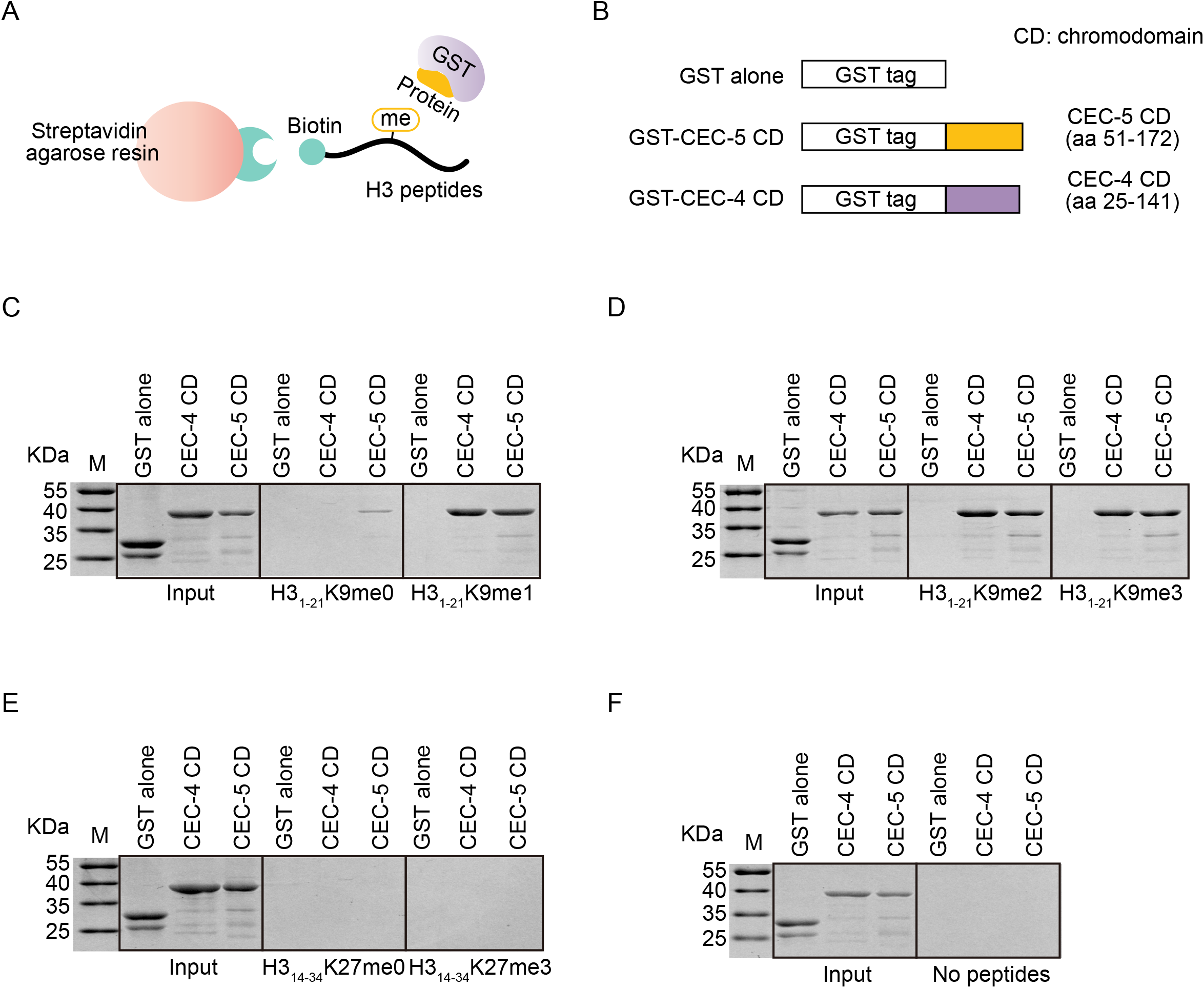
The chromodomain of CEC-5 binds H3K9 peptides in vitro. (A) Schematic diagram of the histone peptide pull-down assay. (B) Schematic diagram of the GST-tagged CEC-5 chromodomain (CEC-5 CD) and CEC-4 chromodomain (CEC-4 CD). (C-F) Coomassie blue-stained SDS[PAGE gels showing binding of the indicated proteins to biotinylated histone H3 peptides. GST alone served as a negative control. The CEC-4 chromodomain was used as a positive control. (C) H3 (1-21) K9me0 (middle) and H3 (1-21) K9me1 (right); (D) H3 (1-21) K9me2 (middle) and H3 (1-21) K9me3 (right); (E) H3 (14-34) K27me0 (middle) and H3 (14-34) K27me3 (right); (F) no peptide (right).

### Forward genetic screening identified that Arg124 in the chromodomain of CEC-5 was required for H3K9me1/2 binding

To further understand the regulation of CEC-5, we performed a forward genetic screening via chemical mutagenesis to search for mutants in which the expression or localization of CEC-5 was altered by clonal screening (Fig. 7A). We isolated a single mutant from approximately one thousand haploid genomes, which disrupted the nuclear puncta formation of CEC-5 (Fig. 7B). We deep sequenced the mutant genome and identified a substitution of Arg124 with cysteine (R124C) in the chromodomain of CEC-5 (Fig. 7C). Notably, Arg124 was conserved in human HP1 (HP1β-CBX1, HP1γ-CBX3, HP1α-CBX5) and PC (PC3-CBX8) proteins, as well as in *C. elegans* CEC-4, CEC-5, and CEC-8, suggesting a critical role of the Arg124 residue (Fig. 7D). In addition, the R124C mutation reshaped the distribution of CEC-5 on heterochromatin and euchromatin (Fig. 7E).

**Figure 7.**
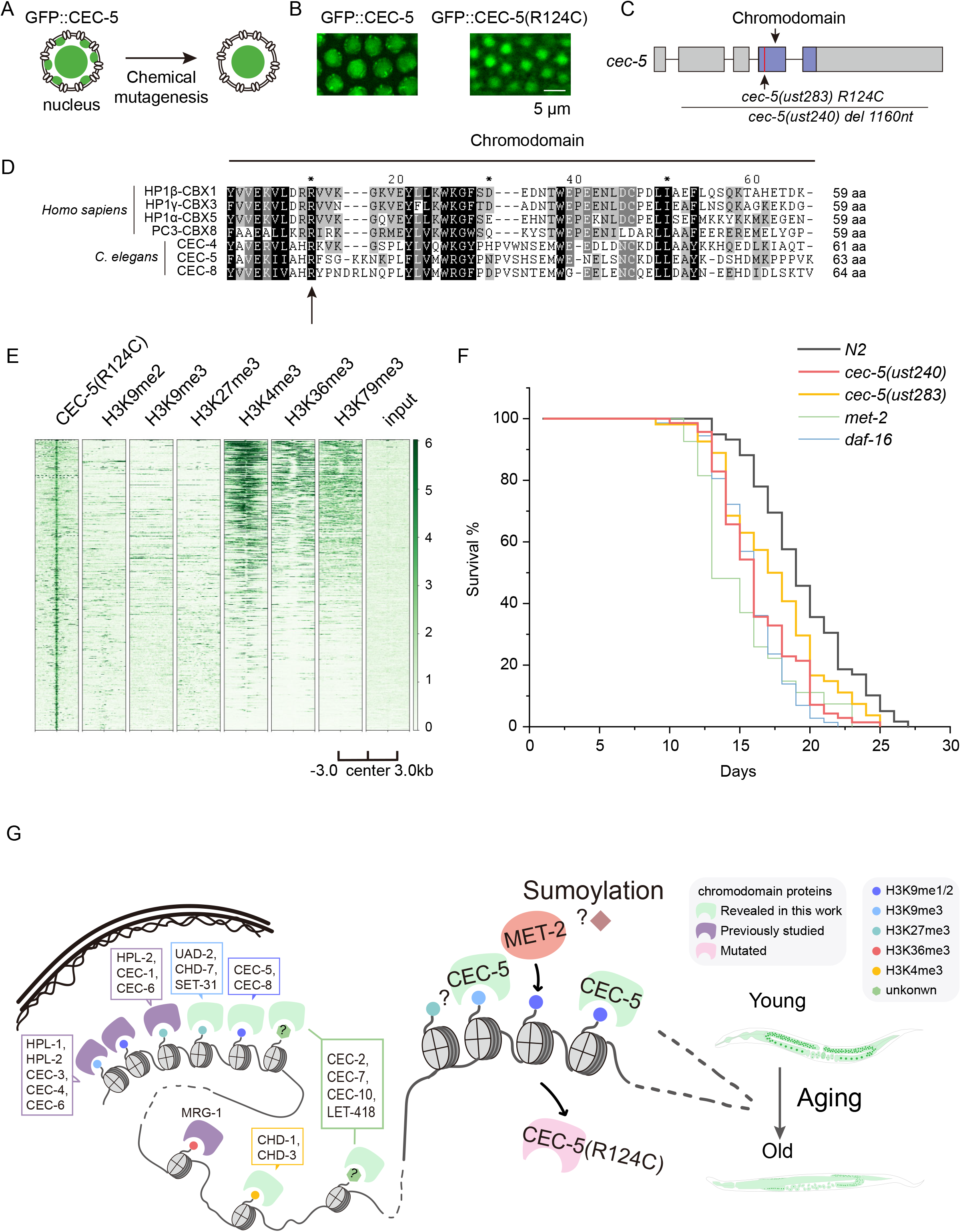
CEC-5 is required for normal lifespan by recognizing H3K9me1/2. (A) Schematic diagram of EMS screening for CEC-5 subcellular delocalization. (B) Images of representative germline nuclei of the indicated adult animals. (C) Schematic diagram of the *cec-5* exon (gray) and chromodomain (purple). (D) ClustalW revealed the multiple sequence alignment of *C. elegans* CEC-4, CEC-5 and CEC-8 chromodomains with different *H. sapiens* HP1 and PC proteins. (E) Heatmap comparing CEC-5(R124C) with heterochromatin histone modifications (H3K9me2, H3K9me3, H3K27me3) and euchromatin histone modifications (H3K4me3, H3K36me3, H3K79me3). (F) Survival curves of the indicated animals. N > 50 worms. (G) A working model summarizing the association of chromodomain proteins with histone modifications in the *C. elegans* genome.

Epigenetic alteration is one of the hallmarks of aging in many organisms (Smeal *et al*. 1996). Aged animals usually show dysregulated repressive heterochromatin (Haithcock *et al*. 2005; Maures *et al*. 2011; Larson *et al*. 2012; Ni *et al*. 2012) (Shumaker *et al*. 2006; Zhang *et al*. 2015; Lee *et al*. 2020). Recent work has reported that the depletion of H3K9me1/2 by *met-2* mutation shortened the lifespan of *C. elegans* (Tian *et al*. 2016). To test whether CEC-5 functions in aging and longevity, we assayed the lifespan of *cec-5* mutants. Both *cec-5(ust240)* and *cec-5(ust283)* were short-lived compared to wild-type N2 animals. Interestingly, *cec-5, met-2* and *daf-16* all exhibited similar shortened lifespans (Fig. 7F).

Taken together, our data suggested that CEC-5, MET-2, and H3K9me1/2 were likely required for the normal lifespan of *C. elegans*. The interaction of CEC-5 occupancy on H3K9me1/2-enriched genomic regions may be critical for CEC-5 function.

## Discussion

In this work, we generated a resource of chromodomain proteins in *C. elegans* by fluorescently tagging the chromodomain proteins using the CRISPR/Cas9 system and performing ChIP-seq, imaging, genetic screening, and functional examination. We presented a functional map of chromodomain proteins (Fig. 7G). We searched for epigenetic factors regulating chromodomain proteins and found that H3K9me1/2 was essential for CEC-5 subcellular localization and heterochromatin association. We conducted a forward genetic screening and identified a conserved residue in the CEC-5 chromodomain required for recognition of H3K9me1/2. *cec-5* is required for the normal life span of *C. elegans*.

### Conservation of chromodomains in eukaryotes

The *C. elegans* genome encodes 21 proteins that contain chromodomains, 9 of which have been reported to have homologs in humans (Table S1). Although the remaining chromodomain proteins in *C. elegans* have no defined homologs in humans, the chromodomain sequences of these proteins still show high similarity to those of human proteins (Table S2) (DasGupta *et al*. 2020). Interestingly, most of these worm-specific chromodomain proteins have a single chromodomain, without any accompanying catalytic domains (Fig. 1B). The complexity of worm-specific chromodomain proteins suggests that flanking sequences or cofactors may modulate the substrate specificity or functions of the proteins. We identified a point mutation in the CEC-5 chromodomain critical for its recognition of methyl-lysine of histone 3. This residue is highly conserved in human HP1 (HP1β-CBX1, HP1γ-CBX3, HP1α-CBX5) and PC (PC3-CBX8) proteins, suggesting a conserved mechanism of H3K9me recognition during evolution.

### Systematically fluorescent tagging of chromodomain proteins

In the last decade, CRISPR/Cas9 technology has revolutionized gene editing to facilitate efficient and precise fluorescent tagging in many organisms. The engineered cells enable examination of protein expression and function in near-native environments. In addition, the transparent and short life-cycled *C. elegans* provides a system to measure the subcellular localization and regulation of proteins under physiological conditions on a life-long scale. The systematic fluorescent tagging of chromodomain proteins provides a resource to investigate the mechanism and function of epigenetic modifications. For example, CEC-5 is localized to the nucleolus and associated with rDNA, suggesting an unknown role of epigenetic modification of histones in rRNA regulation in *C. elegans* (Suka *et al*. 2006; Feng *et al*. 2010; Lee *et al*. 2017).

Using the GFP::CEC-5 strain as a reporter, we found that knocking down the SUMOylation E1 protein *uba-2* or the E2 SUMO-conjugating enzyme *ubc-9* disrupted the nuclear puncta of CEC-5. SUMOylation has been shown to promote the formation and maintenance of silent heterochromatin in yeast and animals (Shin *et al*. 2005; Maison *et al*. 2011; Cubenas-Potts and Matunis 2013; Sheban *et al*. 2022). Our results suggest an underlying mechanism of SUMOylation-controlled heterochromatin regulation.

### The epigenetic regulatory network of chromodomain proteins

In eukaryotic cells, chromosomes are segregated into domains of heterochromatin and euchromatin with distinct functional properties. Heterochromatin is highly condensed, gene poor, rich in transposons or other parasitic genomic elements and generally transcriptionally silent, whereas euchromatin is less condensed, gene-rich, and more easily transcribed (Huisinga *et al*. 2006; Kind and van Steensel 2010; Tamaru 2010). These two silent and active compartments of chromatin differ in characteristic posttranslational modifications (PTMs) of the core histones, as well as in the incorporation of specific histone variants, linker histones, and nonhistone proteins (Towbin *et al*. 2012). Heterochromatin is enriched for H3K27me3 and three methylated states of H3K9, whereas euchromatic domains are marked by H4 acetylation, H3K4me1/2/3, H3K36me1/2/3 and H3K79me1/2/3 (Black and Whetstine 2011; Liu *et al*. 2011). Chromodomain proteins bind both heterochromatin and euchromatin enriched methyl-lysine of histones. The single chromodomain usually recognizes H3K9me2/3 or H3K27me2/3. Paired tandem chromodomains in CHD family proteins are able to bind H3K4me1/2/3. In addition, the chromo barrel domain, a chromo-like domain, can interact with H3K36me2/3.

In this work, we delineated putative histone modifications recognized by chromodomain proteins and related epigenetic factors via systematic analysis of ChIP-seq datasets. Furthermore, by combining genetic screening and in vitro biochemical experiments, we found that CEC-5 binds to H3K9me1/2. Nevertheless, further investigations are required to reveal the direct (or physical) interactions of chromodomain proteins with histones and epigenetic factors. Therefore, a comprehensive interactome (by IP-MS or in vitro pull down assay) of chromodomain proteins will facilitate the understanding of the mechanism and function of chromodomain proteins. GFP-tagged chromodomain proteins could be a promising tool to search for factors modulating heterochromatin.

### Complicate functions of chromodomain proteins

The major function of chromodomain proteins lies in gene expression regulation, genome stability, and three-dimensional genome architecture. In addition to regulating protein coding genes, our work suggested that chromodomain proteins also participate in the control of noncoding sequences. For example, UAD-2 is required for the recruitment of the upstream sequence transcription complex (USTC) to piRNA clusters and promotes piRNA production (Huang *et al*. 2021). HPL-2 and LET-418 are enriched at repetitive elements and prevent aberrant expression of the sequences. Furthermore, nearly all of the tested chromodomain proteins bind to repetitive sequences, implying the general function of chromodomain proteins in genome stability surveillance.

In metazoans, heterochromatic and euchromatic compartments can be distinguished by their spatial organization and association with the nuclear membrane-associated lamina. Recent studies have identified that the perinuclear anchoring of lamina-associated domains in *C. elegans* is facilitated by three chromodomain proteins, CEC-4, MRG-1, and HPL-2 (Gonzalez-Sandoval *et al*. 2015; Cabianca *et al*. 2019; DasGupta *et al*. 2020). Whether other chromodomain proteins participate in the process remains to be determined.

### Chromodomain proteins regulate lifespan and aging

Aging is a complex multifactorial biological process in all living organisms. Among the characterized hallmarks of aging, epigenetic alterations represent a crucial mechanism (Lopez-Otin *et al*. 2013; Sen *et al*. 2016; Zhang *et al*. 2020).

The association between aging and dysregulated repressive heterochromatin has been observed across species, from yeast to humans. In yeast, loss of transcription silencing contributes to aging-related sterility (Smeal *et al*. 1996). In flies and worms, heterochromatin levels positively correlate with lifespan, and aging is associated with deterioration of heterochromatin (Haithcock *et al*. 2005; Maures *et al*. 2011; Larson *et al*. 2012; Ni *et al*. 2012). In mammals, the loss of heterochromatin markers, such as H3K9me, is associated with aging and premature aging diseases (Shumaker *et al*. 2006; Zhang *et al*. 2015; Lee *et al*. 2020). Nevertheless, the aging-associated gain of repressive histone modifications has also been observed. For example, in flies, H3K9me3 levels were found to increase in the aging brain, although they decreased in the aging intestine (Wood *et al*. 2010; Jeon *et al*. 2018). In mice, the levels of H3K27me3 are reduced in senescent fibroblast cells but elevated in the brain of a mouse model of accelerated aging (Bracken *et al*. 2007; Wang *et al*. 2010). In addition, the active transcription mark H3K4me has been shown to be essential for lifespan suppression (Greer *et al*. 2010; Greer *et al*. 2011).

Our data showed that the loss of CEC-5, an H3K9me1/2 reader, and MET-2, the H3K9me1/2 writer, shortens the lifespan of *C. elegans* (Tian *et al*. 2016). Notably, the R124C point mutation on the CEC-5 chromodomain, which disrupted CEC-5 binding to chromatin, also shortened the lifespan of *C. elegans*. How H3K9me1/2 and CEC-5 modulate aging and lifespan requires further investigation.

## Materials and methods

### Strains

Bristol strain N2 was used as the standard wild-type strain. All strains were grown at 20°C unless otherwise specified. The strains used in this study are listed in Table S5.

### Construction of transgenic strains

For chromodomain protein transgenes, endogenous promoter sequences, UTRs, and ORFs of chromodomain genes were PCR-amplified with the primers listed in Table S6. The coding sequence of *gfp::3xflag* and a linker sequence (GGAGGTGGAGGTGGAGCT) (inserted between the ORFs and *gfp::3xflag*) were PCR-amplified with the primers 5’- atggactacaaagaccatgacgg -3’ and 5’- AGCTCCACCTCCACCTCCTTTG -3’. The vector was PCR-amplified from PCFJ151 using the primers 5’- tgtgaaattgttatccgctgg -3’ and 5’- caCACGTGctggcgttacc -3’. Fragments and the vector were fused using a ClonExpress MultiS One Step Cloning Kit (Vazyme C113-02, Nanjing). Chromodomain protein transgenes were integrated into the *C. elegans* genome locus of each gene in situ by using the CRISPR/Cas9 system (Ran *et al*. 2013; Chen *et al*. 2014; Kim and Colaiacovo 2019). The injection mix contained PDD162 (50 ng/µl), plasmid containing chromodomain protein transgenes (50 ng/µl), pCFJ90 (5 ng/µl) and two sgRNAs (30 ng/µl). The mix was injected into young adult N2 animals. The primers used for molecular cloning and sg-RNA sequences are listed in Table S6.

### Microscopy and imaging

Images were collected on a Leica DM4 B microscope. Gonads were dissected in PBS (phosphate-buffered saline) supplemented with 0.2 mM levamisole.

### Chromatin immunoprecipitation (ChIP)

ChIP experiments were performed as previously described (Mao *et al*. 2015). Worm samples in the adult stage were crosslinked in 2% formaldehyde for 30 min. Fixation was quenched with 0.125 M glycine for 5 min at room temperature. Samples were sonicated for 20 cycles (30 sec on and 30 sec off per cycle) at medium output with a Bioruptor 200. The lysates were precleared and immunoprecipitated with 1.5 µL of a rabbit anti-GFP antibody (Abcam, ab290) overnight at 4°C. Chromatin/antibody complexes were recovered with DynabeadsTM Protein A (Invitrogen, 10002D) followed by extensive sequential washes with 150 mM, 500 mM, and 1 M NaCl. Crosslinks were reversed overnight at 65°C. The input DNA was treated with RNase (Roche) for 30 minutes at 65°C, and all DNA samples were purified using a QIAquick PCR purification kit (Qiagen, 28104).

### ChIP-seq

The DNA samples from the ChIP experiments were deep sequenced at Novogene Bioinformatics Technology Co., Ltd. (Tianjin, China). Briefly, 10-300 ng of ChIP DNA was combined with End Repair Mix (Novogene Bioinformatics Technology Co., Ltd. (Tianjin, China)) and incubated for 30 min at 20°C, followed by purification with a QIAquick PCR purification kit (Qiagen). The DNA was then incubated with A-tailing mix for 30 min at 37°C. The 3′-end-adenylated DNA was incubated with the adapter in the ligation mix for 15 min at 20°C. The adapter-ligated DNA was amplified through several rounds of PCR amplification and purified in a 2% agarose gel to recover the target fragments. The average length was analyzed on an Agilent 2100 Bioanalyzer instrument (Agilent DNA 1000 Reagents) and quantified by qPCR (TaqMan probe). The libraries were further amplified on a cBot system to generate clusters on the flow cell and sequenced via a single-end 50 method on a HiSeq1500 system.

### ChIP-seq data downloaded

ChIP-seq datasets of histone modifications, epigenetic factors, and reported chromodomain proteins in *C. elegans* were downloaded from the NCBI GEO or ENCODE databases. The datasets used in this study are listed in Table S3.

### ChIP-seq data analysis

ChIP-seq reads were aligned to the WBcel235 assembly of the *C. elegans* genome using Bowtie2 version 2.3.5.1 (Langmead and Salzberg 2012) by Ben Langmead with the default settings. The SAMtools version 0.1.19 (Li *et al*. 2009) “view” utility was used to convert the alignments to BAM format, and the “sort” utility was used to sort the alignment files. ChIP-seq peaks were called using MACS2 version 2.1.1 (Zhang *et al*. 2008) with subcommand callpeak used with defined parameters (-g ce -B -f BAM -q 0.01 -m 4 50). Deeptools subcommand bamCoverage (version 3.5.0) was used to produce bigWig track for data visualization with defined parameters (--binSize 20 --normalizeUsing BPM --smoothLength 60 --extendReads 150 --centerReads -p 6 2) from bam files. The Integrative Genomics Viewer genome browser (Robinson *et al*. 2011) was applied to visualize signals and peaks. The heatmap was plotted with Deeptools subcommand plotHeatmap (version 3.5.0). The ChIP-seq peaks were annotated with the R package ChIPseeker (Yu et al., 2015).

### Cluster and heatmap analyses

The pipeline of cluster and heatmap analyses was adapted from a previously described method (Griffon *et al*. 2015). Groups of ChIP factors with similar genome-wide signals were determined using the hclust function in R (The R Development Core Team 2009), with pairwise correlation coefficients as the similarity measure. Combined peaks were determined by peaks called from any of the ChIP-seq datasets. The overlapping peaks were merged. The 95^th^ percentile values were extracted from bigWig files using the Python package pyBigWig over each genomic locus of combined peaks. The values were normalized to input signals, logarithm-transformed and then standardized to Z scores. Correlation coefficients were calculated by the cor [Pearson] function in R using the treated values. The agglomeration method “complete” was used.

### Generation of an epigenetic regulatory network

To identify the specific combinations of chromodomain proteins with other epigenetic factors, we generated a colocalization network based on the overlap of binding sites observed in our ChIP-seq datasets using a previously described method (Griffon *et al*. 2015). In brief, we used the IntervalStats tool to compute significant overlaps between binding sites of each pair of factors and calculated the percentage of overlapping sites between the factors. Based on the percentage, we identified strong and moderate colocalization specificity of the factors. The Gephi tool was used to create and visualize the network (Fig. 3B). In the network, the weight of the edges represents how specifically two TFs are associated with each other, whereas the color indicates their percentage of overlapping binding sites. From the network, we identified clusters of strongly correlated chromodomain proteins and epigenetic factors. To highlight these clusters, we further partitioned the network into 8 subnetworks in different colors (Fig. 3B) using an algorithm developed by Blondel *et al*. and implemented in Gephi (Griffon *et al*. 2015). Detailed information on the epigenetic regulatory network is listed in Table S4.

### Candidate-based RNAi screening

RNAi experiments were performed at 20°C by placing synchronized embryos on feeding plates as previously described (68). HT115 bacteria expressing the empty vector L4440 (a gift from A. Fire) were used as controls. Bacterial clones expressing dsRNAs were obtained from the Ahringer RNAi library and sequenced to verify their identity. All feeding RNAi experiments were performed for two generations except for sterile worms, which were RNAi treated for one generation. The genes in the epigenetic RNAi library used in this study are listed in Table S7. Images were collected using a Leica DM4 B microscope.

### Recombinant protein expression and purification

The CEC-4 chromodomain domain (amino acids 25-141) and CEC-5 chromodomain (amino acids 51-172 and 56-182) were PCR amplified, cloned into a plasmid (pET-28a-N8×H-MBP-3C vector), and expressed in *Escherichia coli* Rosetta cells (Novagen). The recombinant proteins were affinity purified through GST tag binding to amylose resin (BioLabs) according to the manufacturer’s instructions.

### Histone peptide pull-down assay

The histone peptide pull-down assay was performed basically as described previously (Huang *et al*. 2021). Briefly, C-terminally biotinylated peptides of *C. elegans* histone H3 (amino acids 1–21 and amino acids 14–34, unmodified or with mono/di/trimethylated lysine, for H3K9me or H3K27me, respectively) were chemically synthesized (SciLight Biotechnology, Beijing, China) and used for the pull-down assay. The peptides were coupled to High-Capacity Streptavidin Agarose Resin (Thermo Scientific) according to the manufacturer’s instructions. Purified GST fusion proteins (3.7 μM) were incubated with the peptide-bead slurry in binding buffer (25 mM Tris-HCl, pH 7.5, 250 mM NaCl, 5% glycerol, 0.1% Triton X-100) for 1 hr at 4°C on a rotator. After washing five times with binding buffer, bound proteins were released from the beads and resolved by SDS□PAGE followed by Coomassie blue staining.

### Lifespan assay

Lifespan assays were performed at 20□. Worm populations were synchronized by placing young adult worms on NGM plates seeded with the *E. coli* strain OP50-1 (unless otherwise noted) for 4–6 hours and then removed. The hatching day was counted as day one for all lifespan measurements. Worms were transferred every other day to new plates to eliminate confounding progeny. Animals were scored as alive or dead every day. Worms were scored as dead if they did not respond to repeated prods with a platinum pick. Worms were censored if they crawled off the plate or died from vulval bursting and bagging. For each lifespan assay, >50 worms were used.

### Statistics

The mean and standard deviation of the results are presented in bar graphs with error bars. All experiments were conducted with independent *C. elegans* animals for the indicated number (N) of times. Statistical analysis was performed with the two-tailed Student’s t test or unpaired Wilcoxon test as indicated.

## Data availability

The raw sequence data reported in this paper have been deposited in the Genome Sequence Archive (Genomics, Proteomics & Bioinformatics 2021) in the National Genomics Data Center (Nucleic Acids Res 2021), China National Center for Bioinformation/Beijing Institute of Genomics, Chinese Academy of Sciences (GSA:), which are publicly accessible at https://bigd.big.ac.cn/gsa/browse/.

## Acknowledgments

We are grateful to the members of the Guang lab for their comments. We are grateful to the International *C. elegans* Gene Knockout Consortium and the National Bioresource Project for providing the strains. Some strains were provided by the CGC, which is funded by the NIH Office of Research Infrastructure Programs (P40 OD010440). This work was supported by grants from the Strategic Priority Research Program of the Chinese Academy of Sciences (XDB39010600), the National Key R&D Program of China (2019YFA0802600), and the National Natural Science Foundation of China (91940303, 31870812, 32070619, 31871300 and 31900434). This study was supported in part by the Fundamental Research Funds for the Central Universities.

## Figure legends of supplementary figures

**Figure S1. The expression of chromodomain proteins**. (A-B) Expression patterns of chromodomain proteins in germline (A) and embryos (B). The expression of a representative image for each chromodomain protein in the indicated tissue or development stage is shown. (C) Brood size of the indicated animals at 20°C. \*\*\**P* < 0.001, \*\**P* < 0.01, \**P* < 0.05; n>20.

**Figure S2. Colocalization of chromodomain proteins with histone modifications identified by ChIP-seq assays**. (A) Distribution of chromodomain protein ChIP-seq peaks on chromosome arms and centers. (B, C, D, E) Heatmap comparing heterochromatin-enriched histone modifications (H3K9me2, H3K9me3, H3K27me3) and euchromatin-enriched histone modifications (H3K4me3, H3K36me3, H3K79me3) with the indicated chromodomain proteins on chromosome arms. (Figure S3 continued)

**Figure S3. Colocalization of chromodomain proteins with histone modifications identified by ChIP-seq assay on chromosome centers**. (A, B, C, D) Heatmap comparing heterochromatin-enriched histone modifications (H3K9me2, H3K9me3, H3K27me3) and euchromatin-enriched histone modifications (H3K4me3, H3K36me3, H3K79me3) with the indicated chromodomain proteins on chromosome centers. (E) A summary of the interaction of chromodomain proteins with histone modifications on chromosome centers.

**Figure S4. The chromodomain proteins mainly bound protein-coding genes**. (A) Gene biotypes bound by chromodomain proteins. (B) GO term analysis of chromodomain protein targets. The area of the bubble reveals the number of genes belonging to the indicated GO categories. Color indicates statistical significance, ranging from highly significant enrichment (red) to enrichment with lower significance (blue). Significantly enriched GO terms were selected based on an adjusted *P* value< 0.05.

**Figure S5. The chromodomain proteins were enriched on transposons and Helitron repetitive sequence families**. (A) Enrichment analysis of chromodomain proteins in repetitive sequence families. The area of the bubble reveals the relative frequency of repetitive sequences belonging to the indicated families. Colors indicate statistical significance, ranging from highly significant enrichment (red) to less significant enrichment (blue). Significantly enriched repetitive sequences were selected based on an adjusted *P* value< 0.05.

**Figure S6. Chromodomain proteins colocalized with histone modifications and other epigenetic regulators**. (A, B, C) The subnetworks showing colocalized factors of the indicated chromodomain proteins. In the subnetworks, nodes indicate individual chromodomain proteins (purple), histone modifications (light pink) and other epigenetic regulators (dark pink); edge colors depict the percentages of overlap between the nodes and weight the colocalization specificity between two factors (see Materials and methods for details). The development stage of animals for each sample is noted as E (embryos), L (larval), and A (adult).

**Figure S7. The subcellular localizations of chromodomain proteins**. (A) Chromodomain proteins localized in the nucleoplasm in embryos. The localization of a representative image for each chromodomain protein is shown (scale bar, 5 μm). The chromosome is marked by mCherry::H2B. (B) Chromodomain proteins localized at nuclear puncta or on chromosomes (scale bar, 5 μm). CEC-5 formed nuclear puncta that colocalized with mCherry::H2B in embryos (left panel); SET-31 colocalized with mCherry::H2B in dividing cells in embryos (right panel, indicated by white arrow). The chromosome is marked by mCherry::H2B. (C) CEC-5 localized to the nucleolus in oocytes (upper panel) and embryos (lower panel) (scale bar, 5 μm). The nucleolus is marked by mCherry::FIB-1. (D) Quantification of the nucleolar localization of CEC-5 and CEC-8 in the indicated cells; N>50. (E) ChIP-seq signals of the indicated chromodomain proteins on 18S and 26S rDNA.

## Supplemental Tables

Table S1. Homologs of chromodomain proteins in humans and *C. elegans*.

Table S2. Chromodomain sequence identity of *C. elegans* chromodomain proteins and human HP1α.

Table S3. ChIP-seq datasets used in this work.

Table S4. Summary of the epigenetic regulatory network.

Table S5. Strains used in this work.

Table S6. sgRNA targeted sequences for CRISPR/Cas9-directed gene editing technology.

Table S7. List of genes used in the candidate-based RNAi screening.

## References

Bannister, A. J., and T. Kouzarides, 2011 Regulation of chromatin by histone modifications. Cell Res 21: 381–395.

Barkan, R., A. J. Zahand, K. Sharabi, A. T. Lamm, N. Feinstein et al., 2012 Ce-emerin and LEM-2: essential roles in Caenorhabditis elegans development, muscle function, and mitosis. Mol Biol Cell 23: 543–552.

Bessler, J. B., E. C. Andersen and A. M. Villeneuve, 2010 Differential localization and independent acquisition of the H3K9me2 and H3K9me3 chromatin modifications in the Caenorhabditis elegans adult germ line. PLoS Genet 6: e1000830.

Black, J. C., and J. R. Whetstine, 2011 Chromatin landscape: methylation beyond transcription. Epigenetics 6: 9–15.

Bracken, A. P., D. Kleine-Kohlbrecher, N. Dietrich, D. Pasini, G. Gargiulo et al., 2007 The Polycomb group proteins bind throughout the INK4A-ARF locus and are disassociated in senescent cells. Genes Dev 21: 525–530.

Broday, L., 2017 The SUMO system in Caenorhabditis elegans development. Int J Dev Biol 61: 159–164.

Cabianca, D. S., C. Munoz-Jimenez, V. Kalck, D. Gaidatzis, J. Padeken et al., 2019 Active chromatin marks drive spatial sequestration of heterochromatin in C. elegans nuclei. Nature 569: 734–739.

Chen, X., F. Xu, C. Zhu, J. Ji, X. Zhou et al., 2014 Dual sgRNA-directed gene knockout using CRISPR/Cas9 technology in Caenorhabditis elegans. Sci Rep 4: 7581.

Coustham, V., C. Bedet, K. Monier, S. Schott, M. Karali et al., 2006 The C. elegans HP1 homologue HPL-2 and the LIN-13 zinc finger protein form a complex implicated in vulval development. Dev Biol 297: 308–322.

Cubenas-Potts, C., and M. J. Matunis, 2013 SUMO: a multifaceted modifier of chromatin structure and function. Dev Cell 24: 1–12.

Cui, M., and M. Han, 2007 Roles of chromatin factors in C. elegans development. WormBook: 1–16.

DasGupta, A., T. L. Lee, C. Li and A. L. Saltzman, 2020 Emerging Roles for Chromo Domain Proteins in Genome Organization and Cell Fate in C. elegans. Front Cell Dev Biol 8: 590195.

Doyon, Y., W. Selleck, W. S. Lane, S. Tan and J. Cote, 2004 Structural and functional conservation of the NuA4 histone acetyltransferase complex from yeast to humans. Mol Cell Biol 24: 1884–1896.

Eissenberg, J. C., 2012 Structural biology of the chromodomain: form and function. Gene 496: 69–78.

Feng, W., M. Yonezawa, J. Ye, T. Jenuwein and I. Grummt, 2010 PHF8 activates transcription of rRNA genes through H3K4me3 binding and H3K9me1/2 demethylation. Nat Struct Mol Biol 17: 445–450.

Flanagan, J. F., L. Z. Mi, M. Chruszcz, M. Cymborowski, K. L. Clines et al., 2005 Double chromodomains cooperate to recognize the methylated histone H3 tail. Nature 438: 1181–1185.

Gonzalez-Sandoval, A., B. D. Towbin, V. Kalck, D. S. Cabianca, D. Gaidatzis et al., 2015 Perinuclear Anchoring of H3K9-Methylated Chromatin Stabilizes Induced Cell Fate in C. elegans Embryos. Cell 163: 1333–1347.

Greer, E. L., T. J. Maures, A. G. Hauswirth, E. M. Green, D. S. Leeman et al., 2010 Members of the H3K4 trimethylation complex regulate lifespan in a germline-dependent manner in C. elegans. Nature 466: 383–387.

Greer, E. L., T. J. Maures, D. Ucar, A. G. Hauswirth, E. Mancini et al., 2011 Transgenerational epigenetic inheritance of longevity in Caenorhabditis elegans. Nature 479: 365–371.

Griffon, A., Q. Barbier, J. Dalino, J. van Helden, S. Spicuglia et al., 2015 Integrative analysis of public ChIP-seq experiments reveals a complex multi-cell regulatory landscape. Nucleic Acids Res 43: e27.

Haithcock, E., Y. Dayani, E. Neufeld, A. J. Zahand, N. Feinstein et al., 2005 Age-related changes of nuclear architecture in Caenorhabditis elegans. Proc Natl Acad Sci U S A 102: 16690–16695.

Harr, J. C., A. Gonzalez-Sandoval and S. M. Gasser, 2016 Histones and histone modifications in perinuclear chromatin anchoring: from yeast to man. EMBO Rep 17: 139–155.

Harrison, M. M., X. Lu and H. R. Horvitz, 2007 LIN-61, one of two Caenorhabditis elegans malignant-brain-tumor-repeat-containing proteins, acts with the DRM and NuRD-like protein complexes in vulval development but not in certain other biological processes. Genetics 176: 255–271.

Huang, X., P. Cheng, C. Weng, Z. Xu, C. Zeng et al., 2021 A chromodomain protein mediates heterochromatin-directed piRNA expression. Proc Natl Acad Sci U S A 118.

Huisinga, K. L., B. Brower-Toland and S. C. Elgin, 2006 The contradictory definitions of heterochromatin: transcription and silencing. Chromosoma 115: 110–122.

Hyun, K., J. Jeon, K. Park and J. Kim, 2017 Writing, erasing and reading histone lysine methylations. Exp Mol Med 49: e324.

Jeon, H. J., Y. S. Kim, J. G. Kim, K. Heo, J. H. Pyo et al., 2018 Effect of heterochromatin stability on intestinal stem cell aging in Drosophila. Mech Ageing Dev 173: 50–60.

Kim, H. M., and M. P. Colaiacovo, 2019 CRISPR-Cas9-Guided Genome Engineering in Caenorhabditis elegans. Curr Protoc Mol Biol 129: e106.

Kind, J., and B. van Steensel, 2010 Genome-nuclear lamina interactions and gene regulation. Curr Opin Cell Biol 22: 320–325.

Koester-Eiserfunke, N., and W. Fischle, 2011 H3K9me2/3 binding of the MBT domain protein LIN-61 is essential for Caenorhabditis elegans vulva development. PLoS Genet 7: e1002017.

Langmead, B., and S. L. Salzberg, 2012 Fast gapped-read alignment with Bowtie 2. Nat Methods 9: 357–359.

Larson, K., S. J. Yan, A. Tsurumi, J. Liu, J. Zhou et al., 2012 Heterochromatin formation promotes longevity and represses ribosomal RNA synthesis. PLoS Genet 8: e1002473.

Lawrence, M., S. Daujat and R. Schneider, 2016 Lateral Thinking: How Histone Modifications Regulate Gene Expression. Trends Genet 32: 42–56.

Lee, D., J. An, Y. U. Park, H. Liaw, R. Woodgate et al., 2017 SHPRH regulates rRNA transcription by recognizing the histone code in an mTOR-dependent manner. Proc Natl Acad Sci U S A 114: E3424–E3433.

Lee, J. H., E. W. Kim, D. L. Croteau and V. A. Bohr, 2020 Heterochromatin: an epigenetic point of view in aging. Exp Mol Med 52: 1466–1474.

Li, H., B. Handsaker, A. Wysoker, T. Fennell, J. Ruan et al., 2009 The Sequence Alignment/Map format and SAMtools. Bioinformatics 25: 2078–2079.

Liu, T., A. Rechtsteiner, T. A. Egelhofer, A. Vielle, I. Latorre et al., 2011 Broad chromosomal domains of histone modification patterns in C. elegans. Genome Res 21: 227–236.

Lomberk, G., L. Wallrath and R. Urrutia, 2006 The Heterochromatin Protein 1 family. Genome Biol 7: 228.

Lopez-Otin, C., M. A. Blasco, L. Partridge, M. Serrano and G. Kroemer, 2013 The hallmarks of aging. Cell 153: 1194–1217.

Maison, C., D. Bailly, D. Roche, R. Montes de Oca, A. V. Probst et al., 2011 SUMOylation promotes de novo targeting of HP1alpha to pericentric heterochromatin. Nat Genet 43: 220–227.

Mao, H., C. Zhu, D. Zong, C. Weng, X. Yang et al., 2015 The Nrde Pathway Mediates Small-RNA-Directed Histone H3 Lysine 27 Trimethylation in Caenorhabditis elegans. Curr Biol 25: 2398–2403.

Maurer-Stroh, S., N. J. Dickens, L. Hughes-Davies, T. Kouzarides, F. Eisenhaber et al., 2003 The Tudor domain ‘Royal Family’: Tudor, plant Agenet, Chromo, PWWP and MBT domains. Trends Biochem Sci 28: 69–74.

Maures, T. J., E. L. Greer, A. G. Hauswirth and A. Brunet, 2011 The H3K27 demethylase UTX-1 regulates C. elegans lifespan in a germline-independent, insulin-dependent manner. Aging Cell 10: 980–990.

McMurchy, A. N., P. Stempor, T. Gaarenstroom, B. Wysolmerski, Y. Dong et al., 2017 A team of heterochromatin factors collaborates with small RNA pathways to combat repetitive elements and germline stress. Elife 6.

Meister, P., B. D. Towbin, B. L. Pike, A. Ponti and S. M. Gasser, 2010 The spatial dynamics of tissue-specific promoters during C. elegans development. Genes Dev 24: 766–782.

Ni, Z., A. Ebata, E. Alipanahiramandi and S. S. Lee, 2012 Two SET domain containing genes link epigenetic changes and aging in Caenorhabditis elegans. Aging Cell 11: 315–325.

Padeken, J., S. Methot, P. Zeller, C. E. Delaney, V. Kalck et al., 2021 Argonaute NRDE-3 and MBT domain protein LIN-61 redundantly recruit an H3K9me3 HMT to prevent embryonic lethality and transposon expression. Genes Dev 35: 82–101.

Perales, R., D. Pagano, G. Wan, B. D. Fields, A. L. Saltzman et al., 2018 Transgenerational Epigenetic Inheritance Is Negatively Regulated by the HERI-1 Chromodomain Protein. Genetics 210: 1287–1299.

Ran, F. A., P. D. Hsu, J. Wright, V. Agarwala, D. A. Scott et al., 2013 Genome engineering using the CRISPR-Cas9 system. Nat Protoc 8: 2281–2308.

Robinson, J. T., H. Thorvaldsdottir, W. Winckler, M. Guttman, E. S. Lander et al., 2011 Integrative genomics viewer. Nat Biotechnol 29: 24–26.

Saltzman, A. L., M. W. Soo, R. Aram and J. T. Lee, 2018 Multiple Histone Methyl-Lysine Readers Ensure Robust Development and Germline Immortality in Caenorhabditis elegans. Genetics 210: 907–923.

Sen, P., P. P. Shah, R. Nativio and S. L. Berger, 2016 Epigenetic Mechanisms of Longevity and Aging. Cell 166: 822–839.

Sheban, D., T. Shani, R. Maor, A. Aguilera-Castrejon, N. Mor et al., 2022 SUMOylation of linker histone H1 drives chromatin condensation and restriction of embryonic cell fate identity. Mol Cell 82: 106–122 e109.

Shin, J. A., E. S. Choi, H. S. Kim, J. C. Y. Ho, F. Z. Watts et al., 2005 SUMO modification is involved in the maintenance of heterochromatin stability in fission yeast. Molecular Cell 19: 817–828.

Shumaker, D. K., T. Dechat, A. Kohlmaier, S. A. Adam, M. R. Bozovsky et al., 2006 Mutant nuclear lamin A leads to progressive alterations of epigenetic control in premature aging. Proc Natl Acad Sci U S A 103: 8703–8708.

Smeal, T., J. Claus, B. Kennedy, F. Cole and L. Guarente, 1996 Loss of transcriptional silencing causes sterility in old mother cells of S. cerevisiae. Cell 84: 633–642.

Suka, N., E. Nakashima, K. Shinmyozu, M. Hidaka and H. Jingami, 2006 The WD40-repeat protein Pwp1p associates in vivo with 25S ribosomal chromatin in a histone H4 tail-dependent manner. Nucleic Acids Res 34: 3555–3567.

Tajul-Arifin, K., R. Teasdale, T. Ravasi, D. A. Hume, J. S. Mattick et al., 2003 Identification and analysis of chromodomain-containing proteins encoded in the mouse transcriptome. Genome Res 13: 1416–1429.

Tamaru, H., 2010 Confining euchromatin/heterochromatin territory: jumonji crosses the line. Genes Dev 24: 1465–1478.

Taverna, S. D., H. Li, A. J. Ruthenburg, C. D. Allis and D. J. Patel, 2007 How chromatin-binding modules interpret histone modifications: lessons from professional pocket pickers. Nat Struct Mol Biol 14: 1025–1040.

Tian, Y., G. Garcia, Q. Bian, K. K. Steffen, L. Joe et al., 2016 Mitochondrial Stress Induces Chromatin Reorganization to Promote Longevity and UPR(mt). Cell 165: 1197–1208.

Towbin, B. D., C. Gonzalez-Aguilera, R. Sack, D. Gaidatzis, V. Kalck et al., 2012 Step-wise methylation of histone H3K9 positions heterochromatin at the nuclear periphery. Cell 150: 934–947.

Vandamme, J., S. Sidoli, L. Mariani, C. Friis, J. Christensen et al., 2015 H3K23me2 is a new heterochromatic mark in Caenorhabditis elegans. Nucleic Acids Res 43: 9694–9710.

Wang, C. M., S. N. Tsai, T. W. Yew, Y. W. Kwan and S. M. Ngai, 2010 Identification of histone methylation multiplicities patterns in the brain of senescence-accelerated prone mouse 8. Biogerontology 11: 87–102.

Weng, C., J. Kosalka, A. C. Berkyurek, P. Stempor, X. Feng et al., 2019 The USTC co-opts an ancient machinery to drive piRNA transcription in C. elegans. Genes Dev 33: 90–102.

Wood, J. G., S. Hillenmeyer, C. Lawrence, C. Chang, S. Hosier et al., 2010 Chromatin remodeling in the aging genome of Drosophila. Aging Cell 9: 971–978.

Wu, X., Z. Shi, M. Cui, M. Han and G. Ruvkun, 2012 Repression of germline RNAi pathways in somatic cells by retinoblastoma pathway chromatin complexes. PLoS Genet 8: e1002542.

Yun, M., J. Wu, J. L. Workman and B. Li, 2011 Readers of histone modifications. Cell Res 21: 564–578.

Zeller, P., J. Padeken, R. van Schendel, V. Kalck, M. Tijsterman et al., 2016 Histone H3K9 methylation is dispensable for Caenorhabditis elegans development but suppresses RNA:DNA hybrid-associated repeat instability. Nat Genet 48: 1385–1395.

Zhang, W., J. Li, K. Suzuki, J. Qu, P. Wang et al., 2015 Aging stem cells. A Werner syndrome stem cell model unveils heterochromatin alterations as a driver of human aging. Science 348: 1160–1163.

Zhang, W., J. Qu, G. H. Liu and J. C. I. Belmonte, 2020 The ageing epigenome and its rejuvenation. Nat Rev Mol Cell Biol 21: 137–150.

Zhang, Y., T. Liu, C. A. Meyer, J. Eeckhoute, D. S. Johnson et al., 2008 Model-based analysis of ChIP-Seq (MACS). Genome Biol 9: R137.

